# Evolutionary innovation through transcription factor promiscuity in microbes is constrained by pre-existing gene regulatory network architecture

**DOI:** 10.1101/2022.07.12.499626

**Authors:** Matthew J. Shepherd, Aidan P. Pierce, Tiffany B. Taylor

## Abstract

The survival of a population during environmental shifts depends on whether the rate of phenotypic adaptation keeps up with the rate of changing conditions. A common way to achieve this is via change to gene regulatory network (GRN) connections – known as rewiring – that facilitate novel interactions and innovation of transcription factors. To understand the success of rapidly adapting organisms, we therefore need to determine the rules that create and constrain opportunities for GRN rewiring.

Here, using an experimental microbial model system we reveal a hierarchy among transcription factors that are rewired to rescue lost function, with alternative rewiring pathways only unmasked after the preferred pathway is eliminated. We identify three key properties - high activation, high expression, and pre-existing low-level affinity for novel target genes – that facilitate transcription factor innovation via gain of functional promiscuity. Ease of acquiring these properties is constrained by pre-existing GRN architecture, which was overcome in our experimental system by both targeted and global network alterations. This work reveals the key properties that determine transcription factor evolvability, and as such, the evolution of GRNs.

## Introduction

During conditions of environmental upheaval and niche transition events, rapid phenotypic adaptation is essential for evolutionary success^1–3^. To understand patterns of diversification in novel environments we need to understand why some evolutionary transitions occur more rapidly than others, and what allows some organisms to succeed where others fail. Key to this is understanding the evolution of gene regulatory networks (GRNs), control circuits common throughout the domains of life that determine the magnitude and timing of gene expression^4^ in response to environmental and internal signals^5, 6^. GRNs are frequent sites of adaptive mutation driving phenotypic evolution^7–9^, and often underscore adaptation to - and survival in - new and changeable environments^10–13^. Determining how these networks evolve and identifying rules governing when and how a regulatory circuit adapts will allow us to better understand their role in determining the evolutionary success of an organism.

Within GRNs, a key mechanism of innovation is transcription factor rewiring, in which transcription factors can gain or lose regulatory connections to target genes creating new network architectures and opportunity for phenotypic innovation^14^. Rewiring events can have dramatic effects on the transcriptome^15^ that can drive phenotypic diversification. However, the majority of past studies on transcription factor rewiring involve retrospective experimental dissection of networks that have already diverged^16–21^, which allows inference of past rewiring events but does not address the evolutionary factors driving the rewiring process. For rewiring to occur, a transcription factor must first have the potential for promiscuous activity (interaction with non-cognate regulatory targets) allowing regulation of a new gene^22^, as is the case for innovation in other proteins^23, 24^. Promiscuity is also referred to as cross-talk, however this term can describe cognate interactions where transcription factors have overlapping regulons, so we avoid it in this study. These non-cognate interactions can become meaningful and drive adaptation if favoured by natural selection^25, 26^.

However, promiscuity cannot be commonplace within GRNs as this will likely result in dysregulation of gene expression and fitness costs for an organism in the environment to which it is adapted^27, 28^.

Whilst there exists some empirical evidence of the importance of transcription factor promiscuity in driving evolutionary innovation^21, 29, 30^, in each case it is unknown why the transcription factor in question became promiscuous as opposed to any of the other regulators within their protein family; what role – if any – GRN structure played in the process; and whether the rewired transcription factors were unique in their ability to become promiscuous within each study system. To address these questions, we set out to investigate the evolution of transcription factor promiscuity using rescue of flagellar motility via rewiring of NtrC documented by Taylor et al. (2015b). In this model system, *Pseudomonas fluorescens* SBW25 is engineered to be non-motile via deletion of the master regulator for flagellar synthesis (*fleQ*) and abolishment of biosurfactant production. Under strong selection for motility, the bacteria rapidly and reliably evolve new regulatory network wiring to rescue flagellar motility (a schematic of these experiments is laid out in Fig.1A)^31^ with the same transcription factor (*ntrC*) repurposed to rescue flagellar driven motility each time, to the exclusion of all other homologous transcription factors within the protein family^32^. However, we do not know the factors that determine this evolutionary preference. To test this, we constructed a double *fleQ ntrC* knockout and placed this mutant under strong selection for swimming motility. This forces evolutionary utilisation of an alternative transcription factor in order to rescue motility, and is an established method for unveiling hidden evolutionary pathways^33^. We reveal a hierarchy among transcription factors that are rewired to rescue lost flagellar function, with alternative rewiring pathways only revealed after the preferred co-opted transcription factor, NtrC, is eliminated. Identification of additional transcription factors capable of gaining promiscuity within the same GRN background allows us to investigate rules governing when and where a transcription factor becomes promiscuous within its GRN, and is important for understanding how these regulatory systems innovate during adaptation to environmental challenges.

**Figure 1:**
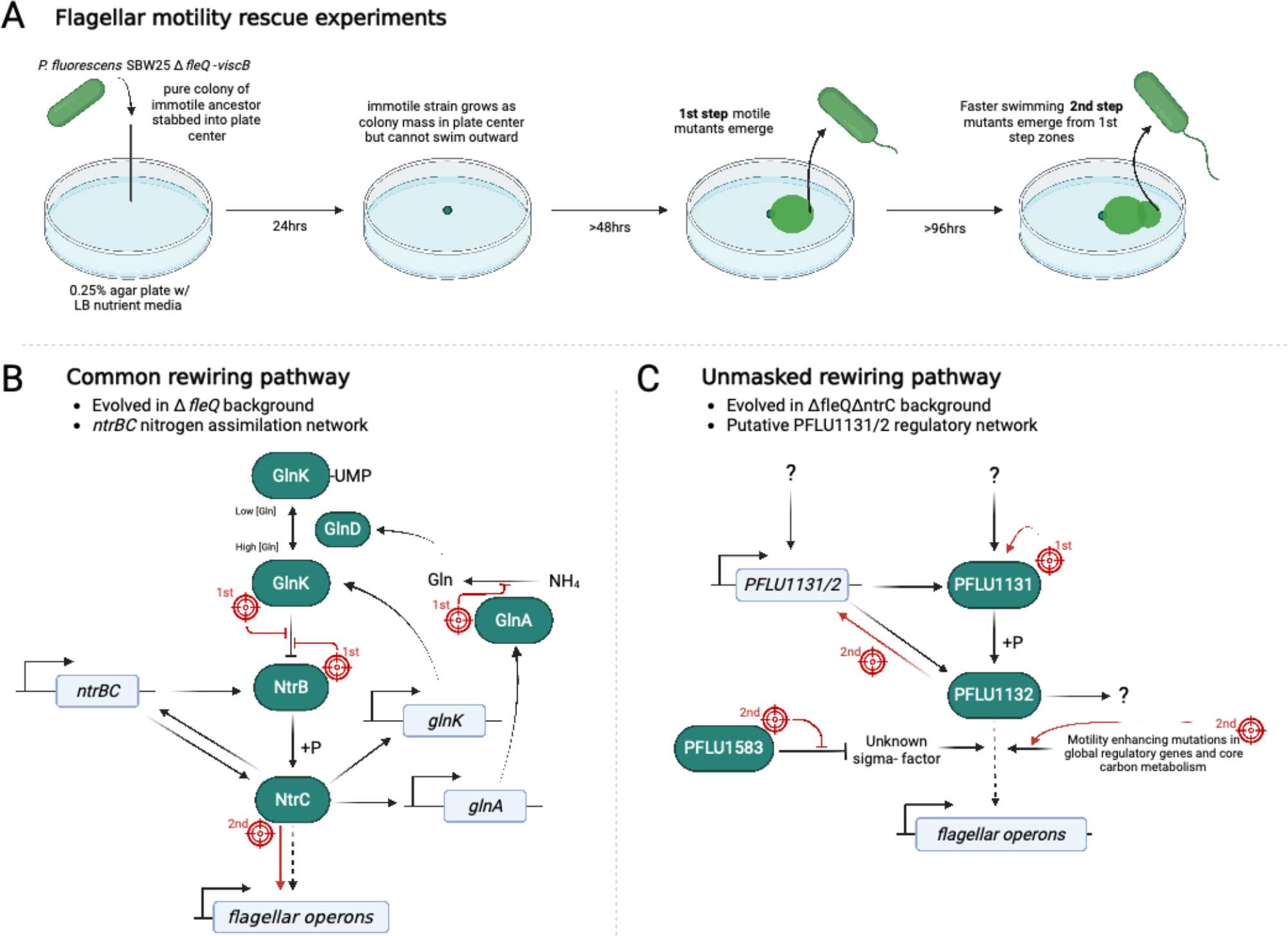
Rewiring of RpoN-EBP transcription factors to rescue flagellar motility through promiscuous control of FleQ-controlled genes. A) Flagellar motility rescue experiment routmap outlines typical progression of motility evolution. Pathway diagrams display key components of the common (B, Taylor et al., 2015) and unmasked (C, this study) rewiring pathways for rescue of flagellar gene expression. Genes are coloured in white, protein components in green. Mutational targets and their effects are shown in red, and whether a mutation occurs in the first or second evolutionary step indicated.

## Results

### Evolutionary rescue of flagellar motility can occur in the absence of *ntrC*, through *de-novo* mutation to an alternative two-component system

Is the reason we only see NtrC co-opted to rescue FleQ function because it is the only transcription factor capable of doing this? It is known that within families of homologous transcription factors there is variation in the ability to bind non-cognate sites^34^, so it is possible that NtrC is unique in its ability to become promiscuous and rewire. FleQ and NtrC are part of a family of 22 structurally related transcription factors called <u>RpoN</u>-dependent <u>e</u>nhancer <u>b</u>inding <u>p</u>roteins (RpoN-EBPs), many of which are predicted to be more structurally similar to FleQ than NtrC^32^. To identify if any other RpoN-EBPs were capable of promiscuity, the double knockout non-motile *P. fluorescens* (Δ*fleQ*Δ*ntrC*) was challenged to rescue flagellar motility in 0.25% agar LB plates. Motile zones re-emerged in a two-step manner (a slow-swimming variant, then a faster-swimming variant) as typical of previous studies^31^. Motile isolates were sampled and whole genome re-sequenced at each step. Motility-granting mutations were identified in the gene *PFLU1131* for all first-step motile isolates (n=15), and in 13/15 cases this was the only mutant gene (Supplementary excel file E1). *PFLU1131* is unstudied and encodes a putative sensor histidine kinase. The gene is situated in an operon between two other genes, *PFLU1130* and *PFLU1132* which encode a hypothetical GNAT-acetyltransferase, and a putative RpoN-dependent transcriptional regulator respectively. PFLU1132 is a known FleQ-homolog^32^ and together with PFLU1131 forms a putative two-component system. The most frequent mutation in *PFLU1131* was an identical 15bp deletion (Fig.2A) in the histidine-kinase phospho- acceptor domain (73%). This mutation results in loss of 5 amino acids (368-GEVAM- 372) in the protein product (henceforth referred to as PFLU1131-del15). Other mutations included a highly similar 15bp deletion resulting in loss of 5 amino acids a few amino acids downstream (369-EVAMG-373), as well as a SNP resulting in A375V. All of these mutations (86% of the first-step mutations) cluster to the same 26bp of the 1770bp *PFLU1131* ORF, and result in amino acid changes in a site directly adjacent to the catalytically active H-box^35^ (amino acids 376-382, Fig.2A) suggesting a significant effect on the catalytic function of the putative kinase. We additionally constructed a *PFLU1132* knockout in the *τιfleQτιntrC* background to identify any further RpoN-EBPs capable of promiscuity. *τιfleQΔntrCΔPFLU1132* failed to rescue flagellar motility within the 6-week assay cut-off (Supplementary excel file E2, n= 192), suggesting evolutionary gain of promiscuity in other transcription factors may not be easily achieved under the conditions we tested.

**Figure 2:**
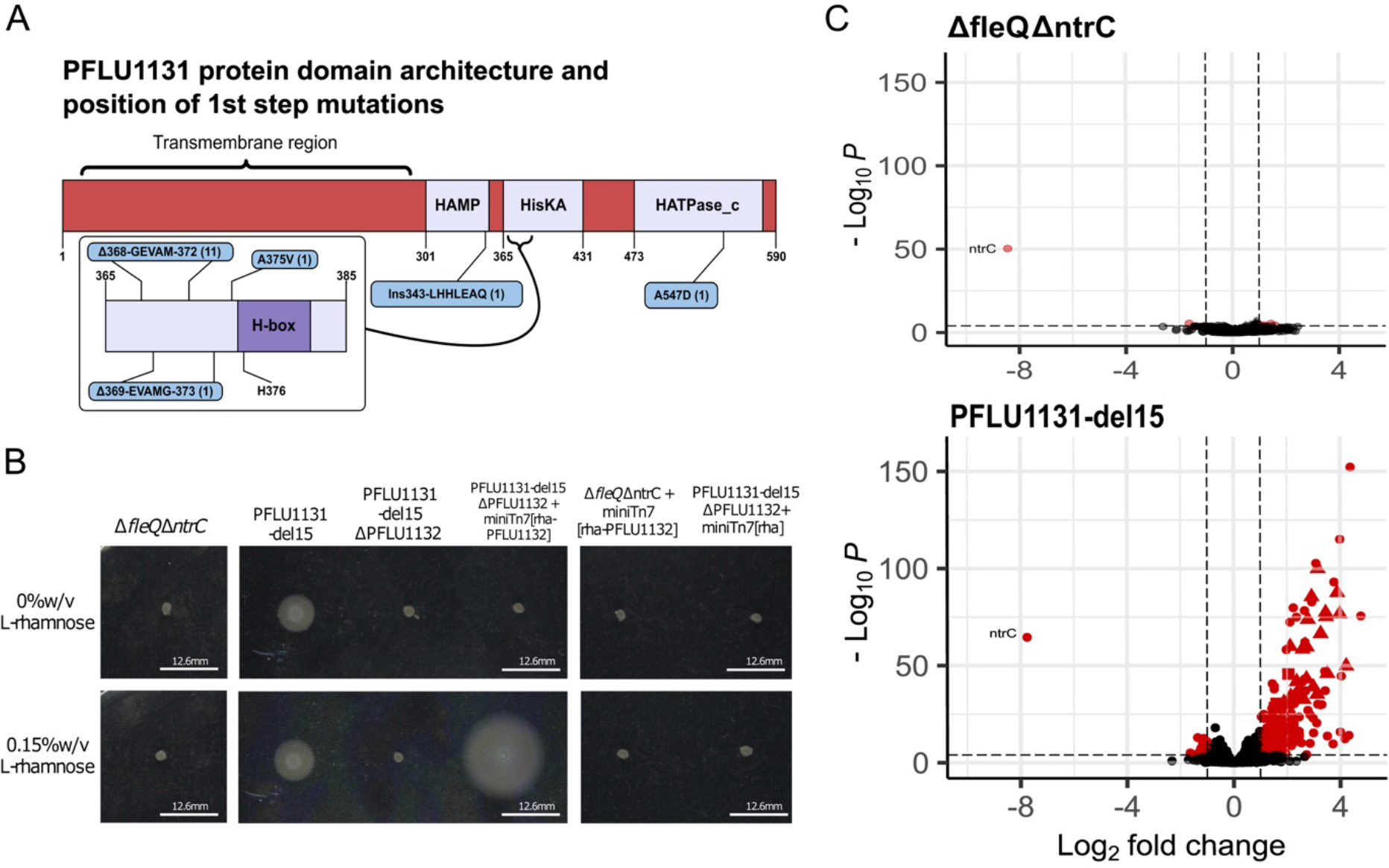
First-step motility rescue mutations in *PFLU1131* A: Diagram of PFLU1131 protein, indicated positions of first-step motility-rescuing mutations. Predicted protein domains (pfam) are indicated in lilac, amino acid positions are indicated as numbers below each domain. Mutations are indicated in blue boxes, with the number of replicate lines gaining each mutation indicated in brackets. **B)** FleQ-homolog encoding gene *PFLU1132* is essential for rescued flagellar motility in the *PFLU1131*-del15 mutant strain and depends on the presence of the kinase mutation. Scale bars (white) = 12.6mm. Transcription factor gene *PFLU1132* is deleted and reintroduced as a single-copy chromosomal insertion expressed from an L-rhamnose inducible promoter system (*rhaSR-PrhaBAD*). The same complementation lacking the *PFLU1131*-del15 mutation, as well as an empty expression system transposon were included as further controls. Photographs of motility after 1 day incubation in 0.25% agar LB plates supplemented with or without 0.15% L-rhamnose for induction of transcription factor expression. **C)** Volcano plots indicating impact of *PFLU1131*-del15 mutation on the transcriptome relative to the *ΔfleQ* ancestor. Red points indicate significantly differentially expressed genes. Triangles indicate flagellar genes, Squares indicate *PFLU1131/2* and adjacent genes, and Circles indicate all other genes.

Together, these results suggest that in the absence of FleQ and NtrC, flagellar motility can be rescued via rewiring of an alternative transcription factor in the same protein family, PFLU1132. To confirm that observed motility phenotypes were dependent on the PFLU1131/2 two-component system, the *PFLU1132* gene was deleted and complemented with and without the presence of the most common first-step kinase mutation (*PFLU1131*-del15). Knockout of *PFLU1132* abolished flagellar motility, and complementation restored motility only in the presence of the kinase mutation (Fig.2B). This experiment was repeated for *ntrBC* and produced the same result (Supplementary Fig.S1). The *PFLU1131*-del15 mutation also grants flagellar motility when *ntrC* is present (i.e., in a Δ*fleQ* background), so does not depend on the absence of *ntrC* to function (Supplementary Fig.S2). Transcriptomic analysis of the *PFLU1131*-del15 mutant by RNA sequencing indicates an alteration to the activity of the PFLU1131/2 two-component system that results in a net up-regulatory effect on the transcriptome (Fig.2C). These regulatory changes include upregulation of the flagellar genes (Supplementary Fig.S4) and the *PFLU1130/1/2* operon. In sum, in the “commonly rewired route”, repeatable mutations in the NtrBC (Fig.1B) two-component system were found to rescue flagellar motility. In the absence of both FleQ and NtrC, we identified the “unmasked rewiring route”, via PFLU1131/2 (Fig.1C), that was also capable of rescuing flagellar motility.

### First-step mutations in the unmasked rewiring pathway grant promiscuous activity to a FleQ homologous transcription factor

In the common rewiring pathway, NtrC, a homolog of FleQ, was recruited to recover lost FleQ function. This occurred as a two-step process; an initial motility-granting mutation to the gene *ntrB* encoding NtrC’s cognate kinase and a secondary motility-enhancing mutation to the helix-turn-helix (HTH) DNA binding domain of the NtrC transcription factor. Similarly, In the Δ*fleQ*Δ*ntrC* background strain, first-step mutations restored slow motility, showing that alternative rewiring routes were possible, but not utilised in the presence of *ntrC*. To determine the mechanism of this unmasked rewiring route we performed whole genome sequencing and transcriptome analysis on ancestral and evolved isolates. In 13/15 first-step mutants, only a single mutation to *PFLU1131* was identified. To confirm that no other mutations were needed for flagellar motility, the *PFLU1131*-del15 mutation was introduced into the ancestor (Δ*fleQ*Δ*ntrC*). The resulting engineered strain was motile (Supplementary Fig.S2). As no mutational change to the DNA binding domain of PFLU1132 is needed, this suggests a non-specific mechanism through which this transcription factor induces the flagellar genes.

To investigate this, our transcriptomic data was used to assess the impact of the *PFLU1131*-del15 mutation on the expression of a list of genes^36^ controlled by all 22 homologous transcription factors (part of the RpoN-EBP family) present in *P. fluorescens* SBW25^37^. Regulatory changes predictably include upregulation of the flagellar genes (Supplementary Fig.S3) and the *PFLU1130/1/2* operon, but also include upregulation of many other genes. In particular, we saw significant upregulation for 54% of all RpoN-EBP controlled genes in the unmasked rewiring pathway (Fig.3). For comparison, 70% of RpoN-EBP controlled genes were upregulated in the common rewiring pathway (via NtrC)^34^. Flagellar genes account for 12% of all RpoN-EBP controlled genes, and whilst the native regulatory targets of PFLU1132 are unknown it is unlikely that it natively regulates 54% of these.

**Figure 3:**
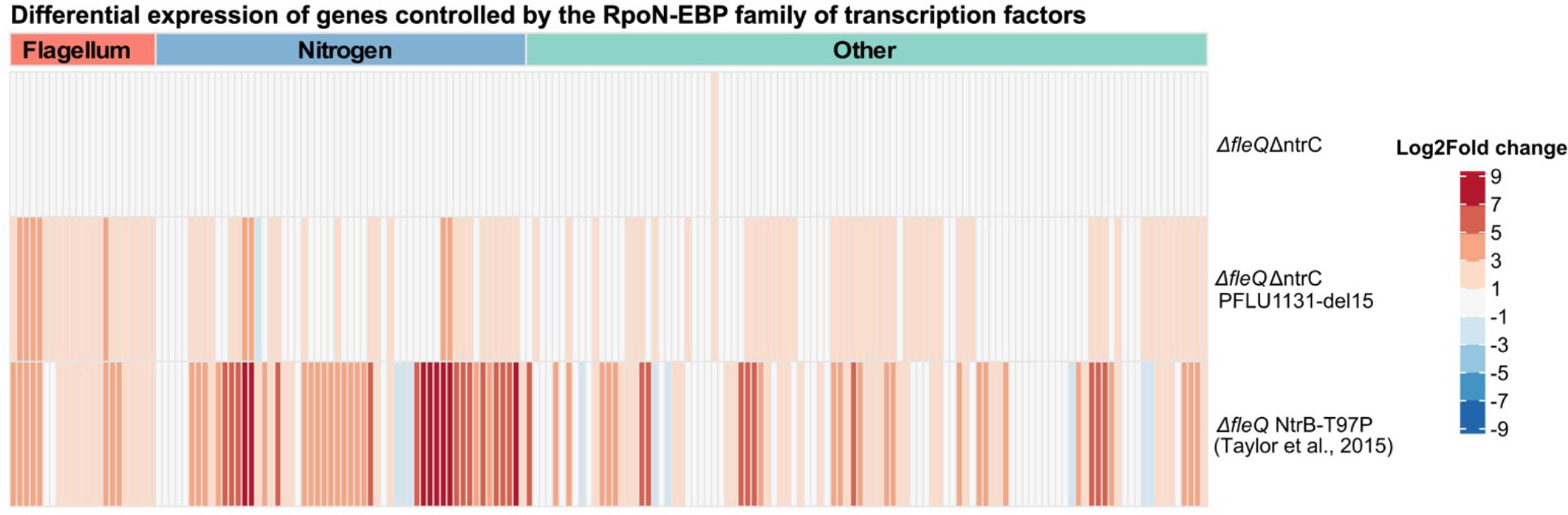
Log2Fold changes in gene expression for RpoN-EBP controlled genes after first-step motility mutations, relative to. Δ*fleQ* **ancestor.** Δ*fleQ*Δ*ntrC* and Δ*fleQ*Δ*ntrC* first-step differentially expressed genes were determined by RNA sequencing (this work). Δ*fleQ* first-step differentially expressed genes were identified by RNA microarray and originally reported by Taylor *et al.* 2015. List of RpoN-EBP controlled genes was obtained from Jones *et al.,* 2007. Groups of genes by function are indicated by the coloured bars above the plot: Red – Flagellum, Blue – Nitrogen, Teal – Other/unknown.

Similarly, the first step motile mutant reported in Taylor et al., (2015b) leads to upregulation of genes known to be involved in nitrogen assimilation (as expected as mutations are located in the Ntr pathway), however it also results in upregulation of many genes with no known regulatory link to NtrC. The large percentages of the RpoN regulon induced by these specific kinase mutations, in both the common and unmasked rewiring pathways, suggest first-step mutations confer a state of promiscuous regulatory activity to PFLU1132 resulting in non-cognate interactions across many RpoN-EBP controlled genes including the flagellar genes.

### Evolutionary rescue of motility through unmasked rewiring pathway is significantly constrained

If alternative rewiring routes are available to natural selection – why have we not seen them utilised in the presence of the commonly rewired route? In the unmasked rewiring pathway, we considered the time taken for swimming mutants to evolve and the strength of the evolved phenotype in comparison to the common rewiring route. Time to emergence (the length of time taken for motility to evolve on the soft agar plate) was recorded for all replicate experimental lines of the Δ*fleQ*Δ*ntrC* ancestor, as well as Δ*fleQ* as a comparison. Strikingly, only 9.4% of replicate experimental lines (n=160) of Δ*fleQ*Δ*ntrC* evolved within the 6-week assay cut-off compared to 100% for the Δ*fleQ* background (n=22). Initial first-step motile Δ*fleQ*Δ*ntrC* strains also evolved within an average of 18.7 days from the assay start (Fig.4A), significantly longer than Δ*fleQ* took to rescue motility (4.2 days, P = 0.002, Dunn test). After first-step PFLU1131 mutations, second-step mutants evolved rapidly in Δ*fleQ*Δ*ntrC* lines within an average of 2.2 days (Fig.4A). This was faster than the second-step mutants in the Δ*fleQ* background which took 3.3 days to emerge (*P* = 0.0017, Dunn test).

**Figure 4:**
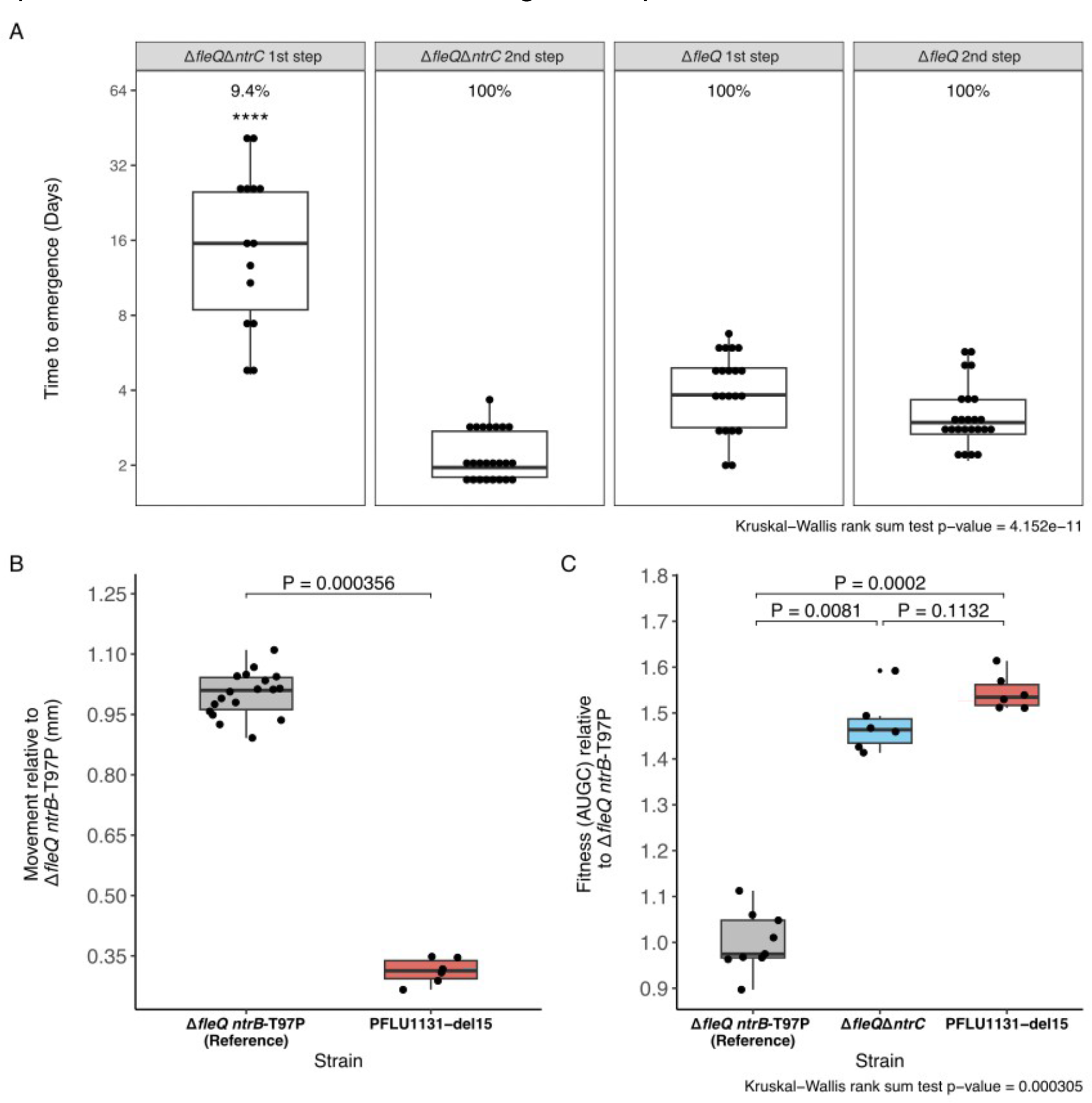
Phenotypes of motility rescue mutants. A) Time to emergence of motile zone (Days) for first-and second-step mutants in the Δ*fleQ*Δ*ntrC* and Δ*fleQ* backgrounds. Percentages above each plot indicate proportion of replicate evolving that step within 6 week assay cutoff. **** indicates significant difference in mean time to emergence from all other groups (Dunn test). **B)** Race assay as measure of motility fitness. Distance moved over 24hrs in 0.25% Agar LB plates measured relative to the Δ*fleQ ntrB*-T97P mutant. P-value indicated is generated from a two- sample Wilcox test. **C)** Fitness in LB broth measured as area under the 24hr growth curve (AUGC) relative to Δ*fleQ ntrB*-T97P. P-values above plots generated from Dunn tests. AUGC values and growth curve plots are provided in Supplementary Fig.S3. For all boxplots – box represents first to third quartile range, middle line represents median value, whiskers range from quartiles to maxima and minima.

The greatly increased time, and low frequency for rescue motility in an Δ*fleQ*Δ*ntrC* background may be reflective of a small pool of accessible PFLU1131 mutations that can grant promiscuity to PFLU1132, perhaps due to a small mutational target size.

One replicate line gained a *de novo* mutation in the DNA mismatch-repair gene *mutS* (Supplementary excel file E1), resulting in a frame shift and probable loss of function^38^. This strain, along with strains derived from it possessed large numbers of additional SNPs (>80) including a SNP in PFLU1131. Loss of function to *mutS* is known to enhance mutability in *Pseudomonads*^39^, and its occurrence may increase access to motility-rescuing mutations in PFLU1131.

From a phenotypic perspective, the first-step mutations in the *PFLU1131* pathway in a Δ*fleQ*Δ*ntrC* strain, provide a far poorer motility phenotype than the analogous first-step mutations in a *ΔfleQ* background (Fig.4B). In a race assay, *PFLU1131*-del15 mutants swam 0.31 mm for every 1 mm swam by the most common first step mutant in the common rewiring route (P=0.000356, Wilcox test). One possible explanation for this poor motility is a lack of significant upregulation for the flagellar filament subunit FliC (Supplementary Fig.S4) in the *PFLU1131*-del15 mutant.

The *PFLU1131*-del15 mutant additionally does not confer a significant defect to growth in shaking LB broth compared to its ancestral strain (Fig.4C, average relative area under the growth curve (rAUGC) of 1.55 and 1.48 respectively, P=0.1132, Dunn test), and grew significantly better than the most common first step mutant in the common rewiring route (rAUGC of 1, P=0.0081, Dunn test). The poor motility phenotype and the lack of significant fitness cost associated with *PFLU1131*-del15 likely indicates that the PFLU1131/2 system is a weaker activator of the flagellar motility that incurs less sever pleiotropy compared to NtrBC.

These findings indicate that the unmasked pathway is the far poorer option for gaining promiscuous regulation of the flagellar genes compared to the common NtrBC pathway - motility rescue in the τι*fleQ*τι*ntrC* background took significantly longer to occur, was significantly less frequent, and provides a significantly poorer flagellar motility phenotype with lower pleiotropic fitness cost. However, it is not clear why this should be the case – both NtrC and PFLU1132 are FleQ-homologs, and are gaining mutations to their cognate kinases that unlock promiscuity. This suggests that there are other factors that constrain the evolution of promiscuity in PFLU1132.

### Motility enhancing second-step mutations suggest alterations to global gene regulatory network can facilitate promiscuity in unmasked rewiring pathway

Following the first-step mutations in PFLU1131 that unlock flagellar motility, our motile isolates develop second-step mutations that act to boost motility speed (Supplementary Fig.S5A), often accompanied by significant pleiotropic fitness costs for growth in LB broth (Fig. S5B). These second-step mutations offer clues to the nature of constraining factors limiting innovation through promiscuity of PFLU1132, as they represent evolutionary solutions to the poor motility provided to the first-step mutations.

Whole genome resequencing identified a diverse set of motility enhancing second-step mutations occurring at both a local (PFLU1131/2 locus) and global regulatory scale (Fig.5A, full details of all mutations provided in supplementary excel file E1, second step n = 18 (several first step lineages generates multiple second step zones). In contrast to the frequent second-step mutations observed in the HTH DNA-binding domain of NtrC^31^, no analogous mutations were observed in the same domain of PFLU1132. Mutations that did occur in the *PFLU1132* gene were parallel and identified in only 2 lines, impacting the receiver domain of the transcription factor, which may further increase activity of the PFLU1132 regulator by modifying the interactions with its kinase. One line gained a secondary *PFLU1131* mutation, a SNP in the HATPase domain along with the first-step SNP A547D already present (Fig.2B), which may further boost kinase interactions. A mutation to the promoter region of the PFLU1131/2 operon was also observed.

**Figure 5:**
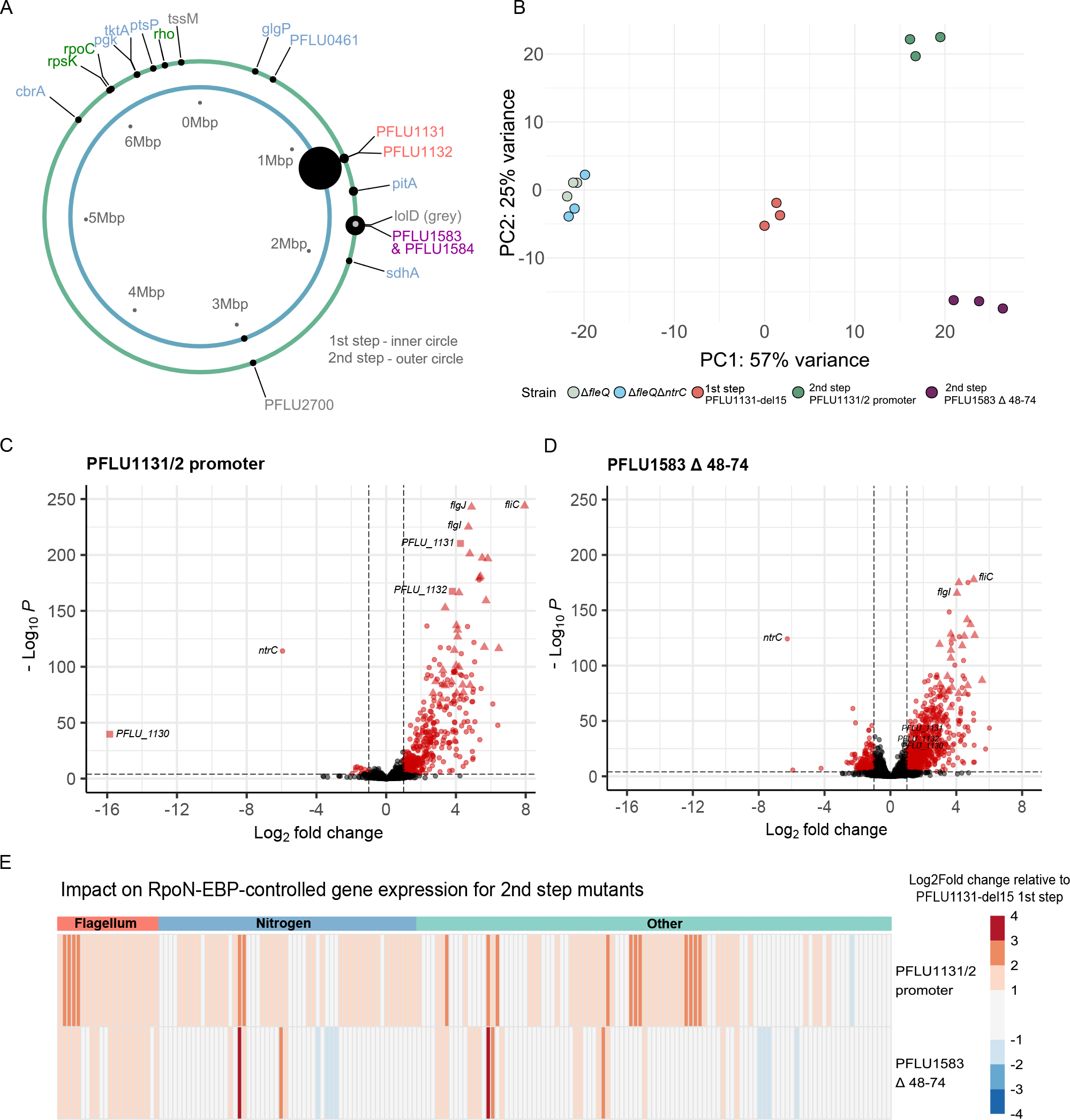
Diverse second-step mutations indicate multiple global regulatory strategies for enhancing promiscuity through PFLU1132. A) Diagram of mutational distribution across the SBW25 chromosome for first and second step motility mutants. Each ring represents the chromosome, with black dots indicating non-synonymous mutations. The size of each dot is proportional to the number of mutations occurring in that locus in independent replicates. Mutational target names are coloured by functional category: Red – *PFLU1131/2*, Purple – *PFLU1583/4*, Blue – Metabolic, Green – Global regulatory, Grey – other. **B)** Principle component analysis of transcriptomic data for ancestral, first-step and second-step motility mutants. **C and D)** Volcano plots of differentially expressed genes for second-step mutations relative to Δ*fleQ* ancestor. Red points are significantly differentially expressed. Triangles indicate flagellar genes, Squares indicate *PFLU1131/2* and adjacent genes, and Circles indicate all other genes. **E)** Log2Fold change in gene expression for RpoN-EBP controlled genes after *PFLU1583* Δ48-74 mutation – relative to the first-step *PFLU1131*-del15 mutant. RpoN-EBP controlled genes grouped by function are indicated by the coloured bars above the plot: Red – Flagellum, Blue – Nitrogen, Teal – Other/unknown

Other secondary mutations can be grouped into 3 other broad categories. (i) The first are mutations in the operon *PFLU1583*/*4*, accounting for 24% of secondary mutations (Fig.5A, Purple). This gene pair encodes a putative anti-sigma factor PP2C-like phosphatase and a putative STAS family anti-sigma factor antagonist (*PFLU1583* and *PFLU1584* respectively). Mutations in *PFLU1583* were generally loss of function suggesting loss of repression on an unknown sigma factor. This may act on RpoN the partner sigma factor of PFLU1132 in regulating gene expression, however this cannot be tested easily through *rpoN* knockout, and *P. fluorescens* SBW25 encodes ∼31 other putative sigma factors^37^ which may instead be the mechanistic targets of this mutation. Transcriptomic analysis of a *PFLU1583* mutant (PFLU1583Δ48-74) identified sigma factor RpoE as upregulated, however *rpoE* knockout did not negate the motility enhancing effect of the mutation (Supplementary Fig.S6A). To test that PFLU1583/4 are not acting to enhance motility through another RpoN-EBP, we engineered *PFLU1583*Δ48-74 in a Λ1*fleQ*Λ1*ntrC* background in the absence of a *PFLU1131* mutation. This strain was immotile, indicating that these anti-sigma factor mutations depend on the function of the PFLU1131/2 system to confer a swimming phenotype (Supplementary Fig.S6B). (ii) The second category of mutations impact core gene expression components that will affect global GRN function (Fig.5A, Green). These included mutations to *rho*, *rpsK* and *rpoC* that encode core gene expression machinery and will impact expression of most genes in the genome. (iii) The final category of motility enhancing mutations fit a metabolic theme (Fig.5A, blue). These mutations occur in genes involved in central carbon metabolism, which will likely have a significant impact on global gene expression through alterations to central carbon flux and may act on RpoN indirectly – for example via core carbon metabolic regulator CbrB (an RpoN-EBP) – or signal to the PFLU1131 sensor-kinase via internal metabolic flux.

The diversity and nature of these second-step mutations suggest global regulatory strategies to facilitate flagellar gene expression through the unmasked rewiring pathway. To understand their regulatory impact, transcriptome analysis was performed on a pair of representative mutants: the most common second-step mutant (anti-sigma factor mutant, *PFLU1583* Δ48-74) which represented a mutation with global regulatory effects, and the *PFLU1131/2* promoter mutation as a representative of mutation with predicted “local” regulatory effects. Together these mutations represent the two loci that constitute 56% of identified second-step mutations.

Principle component analysis indicates that the transcriptomic profile of the anti-sigma factor mutant differs significantly from the profile of the *PFLU1131/2* promoter mutation. Both mutations resulted in similar variation across PC1 relative to the first-step *PFLU1131*-del15 mutant, but opposite variation in PC2 (Fig.5B). Both the *PFLU1131/2* promoter and anti-sigma factor mutations result in net up-regulatory effects on the transcriptome reflected by positive skewed volcano plots (Fig.5C and Fig.5D) albeit with differing expression patterns. Both mutations have the effect of further upregulating RpoN-EBP controlled genes with 22% and 60% being upregulated for the anti-sigma factor mutant and the *PFLU1131/2* promoter mutant respectively (Fig.5E).

In general, for the unmasked rewiring pathway, genomic and transcriptomic analysis of second-step mutations indicate that second-step mutations are increasing promiscuous activity of the rewired transcription factor (PFLU1132), albeit through differing (local and global) mechanisms. These results highlight that diverse mutations with both targeted and global regulatory effects can influence promiscuity to facilitate rewiring and innovation in a transcription factor.

### Second-step promoter capture mutation suggests importance of transcription factor expression in gaining promiscuity

Although second-step mutations were diverse in the unmasked rewiring pathway, one promoter capture event in particular provides evidence for the role of increased expression of the rewired transcription factor in strengthening the motility phenotype. For promiscuous activation of non-cognate genes, the PFLU1132 transcription factor will need to first saturate its native regulatory interactions before it can engage in low-affinity promiscuous interactions with FleQ-controlled genes. High expression of active transcription factor can provide these conditions, elevating the concentration of transcription factor in the cell to permit more frequent promiscuous action^40, 41^.

Whilst many of the second-step mutations detailed above may impact *PFLU1131/2* expression indirectly (RpoC, Rho), the *PFLU1131/2* promoter mutation directly effects expression of the two-component system. This mutation is a 1.59Kbp deletion resulting in total loss of the *PFLU1130* gene, and in *PFLU1131/2* becoming part of the *PFLU1127/8/9* operon (Fig.6A). Upstream of this new combined operon sits a predicted RpoN binding site^36^, so this deletion positions *PFLU1132* in a new operon under the control of RpoN which may create a positive feedback loop where this RpoN-dependent regulator drives its own expression – either through promiscuous regulation or native control of this promoter. This genetic rearrangement significantly upregulates both *PFLU1131* and *PFLU1132* with 4.26 and 3.79 Log2Fold increases in expression compared to the non-motile Δ*fleQ* strain respectively (Fig.6B, raw values for this and other tested strains provided in supplementary excel file E5).

**Figure 6:**
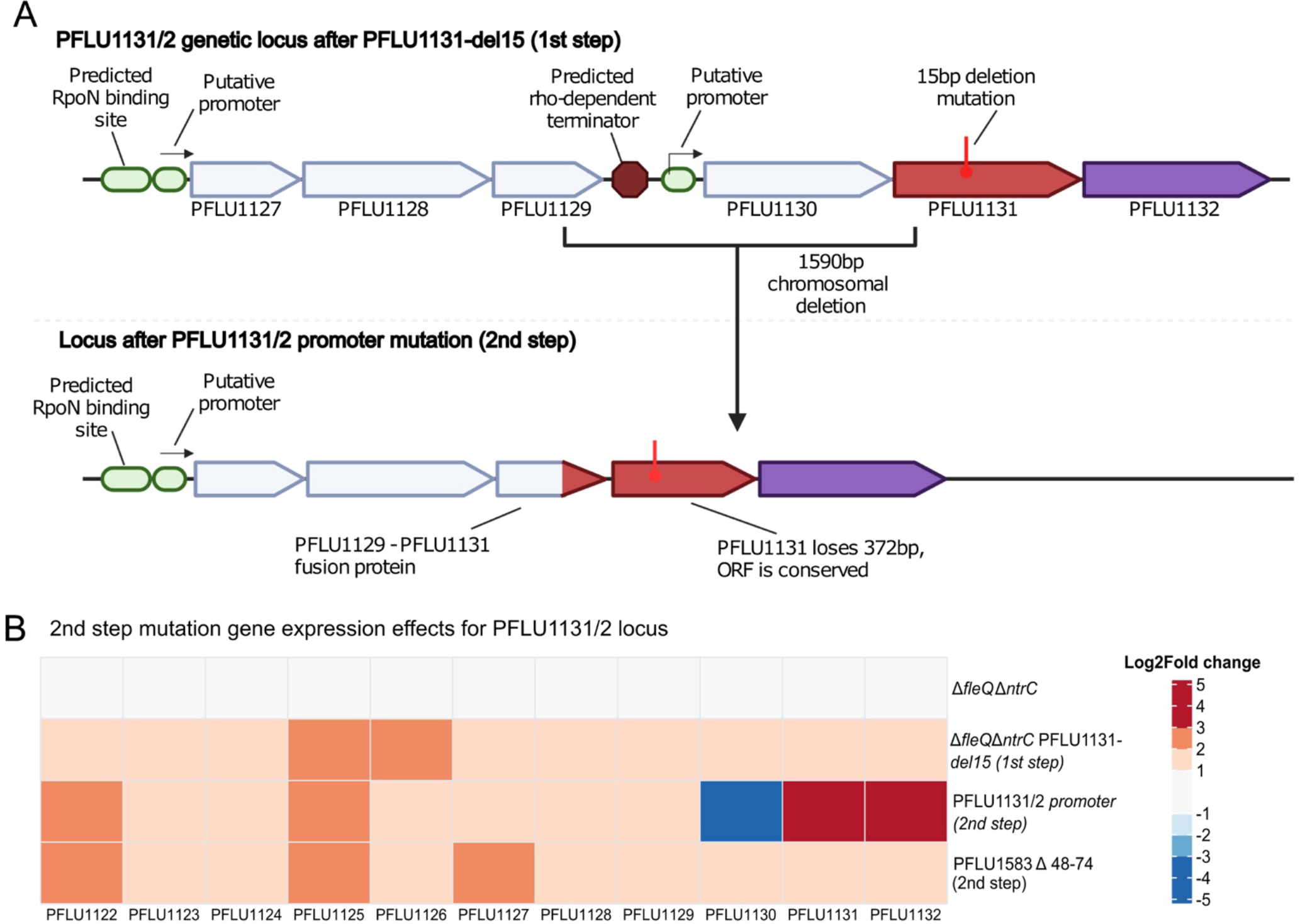
Second-step *PFLU1131/2* promoter mutation increases *PFLU1131/2* expression. A) *PFLU1130/1/2* genetic locus in the SBW25 chromosome, and impact of the 1.59kbp deletion identified in one second-step motile mutant. Location of the RpoN binding site upstream of PFLU1127/8/9 was predicted by Jones et al., (2007). The presence of the rho-dependent terminator was determined by use of RhoTermPredict^71^ **B)** Log2Fold change in gene expression (relative to Δ*fleQ* ancestor) heatmap of *PFLU1130/1/2* and nearby RpoN-EBP controlled genes, including *PFLU1127/8/9* in the ancestral Δ*fleQ*Δ*ntrC* strain, first and second step motility mutants.

Upregulation of this two-component system corresponds with the 60% increase in expression of RpoN-EBP controlled genes discussed above (Fig.5E), indicating an increase in promiscuous transcription factor activity. The 1.59Kbp deletion results in loss of an intergenic region that typically separates the two combined operons. This region is predicted to contain a rho-dependent terminator (Supplementary Excel file E3) which may typically prevent transcriptional readthrough into *PFLU1130/1/2*.

Such a terminator will restrict *PFLU1131/2* expression via readthrough and constrain its ability to achieve high concentrations that may facilitate promiscuity. The *rho* mutation observed in another second-step motile strain may also act to increase readthrough at this site. In comparison to this promoter mutation, the other second- step mutation tested (anti-sigma factor *PFLU1583* Δ48-74) does not significantly impact *PFLU1131/2* expression (Fig.6B), so this mutation likely influences the promiscuity of the *PFLU1132* transcription factor through a separate mechanism.

In summary, expression of *PFLU113/2* appears to be a major factor constraining promiscuous regulatory activity and motility rescue through this regulatory system, and overcoming this constraint facilitates evolutionary innovation through this unmasked rewiring route.

### Increasing transcription factor expression in motile mutants results in faster motility phenotypes

To test the importance of transcription factor gene expression in gain of promiscuous activity, we made use of the complementation strains presented in Fig.2C and Supplementary Fig.S1. These experiments produced strains where the common (*ntrC)* and unmasked (*PFLU1132)* rewired transcription factor genes are deleted from their native loci and reintroduced on an L-rhamnose titratable promoter system. This was done in the presence of their respective kinase mutations that conferred first-step slow spreading motility in each rewiring pathway. Concentration of L-rhamnose added to the media positively correlated with distance moved (Fig.7A) for both the common rewiring pathway (i.e., NtrC expression system; ρ = 0.984, Spearman’s test) and the unmasked rewiring pathway (i.e., PFLU1132 expression system; ρ = 0.865, Spearman’s test). We confirmed increasing L-rhamnose concentration increased expression from the L-rhamnose titratable system (RhaSR-PrhaBAD) in our bacterial strain backgrounds by testing LacZ activity with increasing L-rhamnose concentration for *lacZ* under control of the L-rhamnose titratable promoter construct in the *ΔfleQ* and *ΔfleQ*Δ*ntrC* backgrounds. In both cases LacZ activity positively correlated with L-rhamnose concentration (ρ = 0.981 and ρ = 0.955 respectively, Spearman’s tests), indicating that increasing L-rhamnose concentration results in increasing expression of the gene introduced in the *rhaSR-PrhaBAD* expression system (Supplementary Fig.S7).

**Figure 7:**
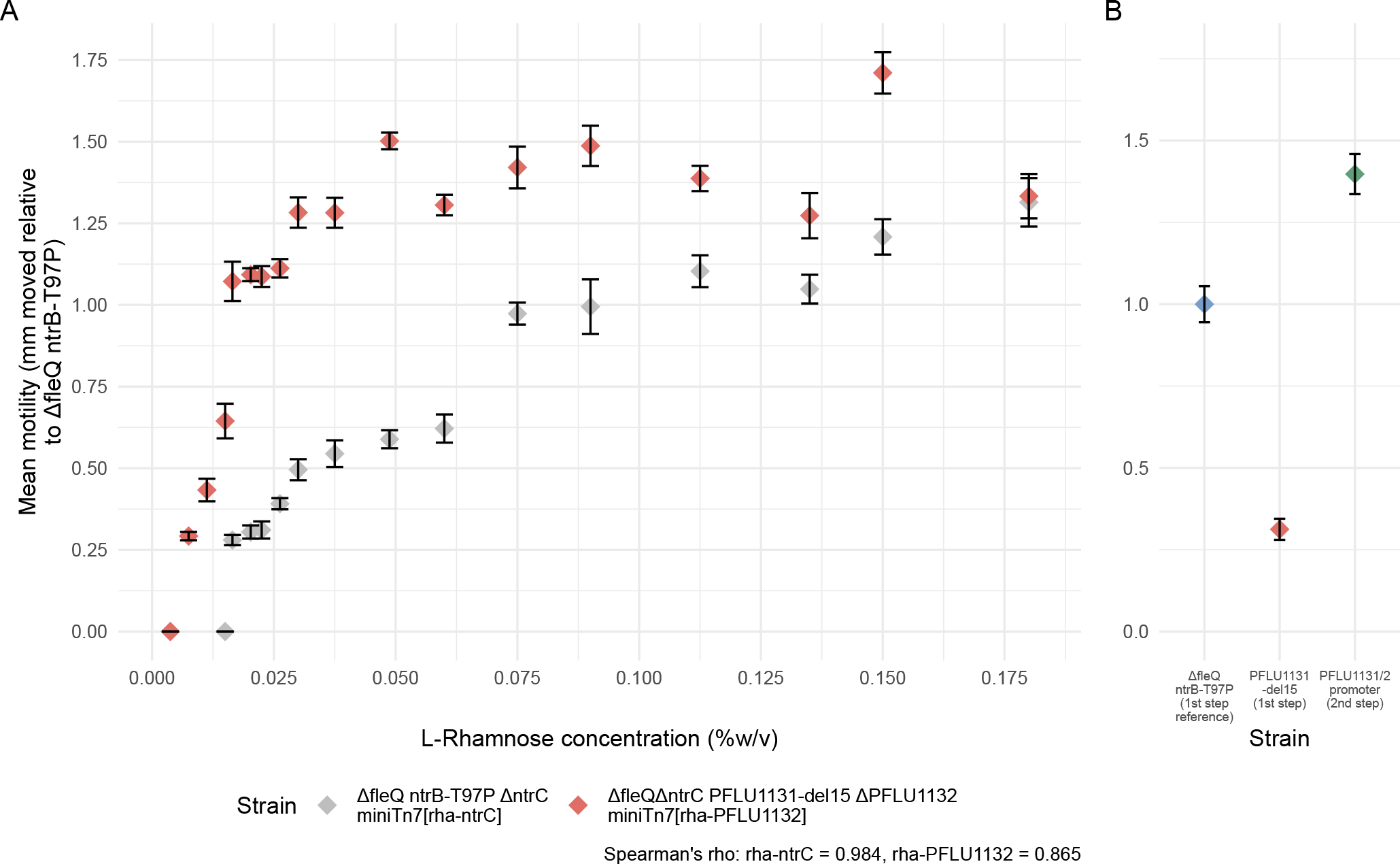
Impact of RpoN-EBP gene expression on motility speed. A) Mean motility speed (mm relative to Δ*fleQ ntrB*-T97P) plotted against increasing L- Rhamnose concentration (%w/v). Mean of six biological replicates for each point. first-step motility mutants *ΔfleQ ntrB*-T97P and Δ*fleQ*Δ*ntrC PFLU1131*-del15 had their respective RpoN-EBPs knockout out, and reintroduced on a miniTn7 transposon with the RpoN-EBP under control of the PrhaBAD and *rhaSR* rhamnose inducible expression system. In full, these were – Δ*fleQ* ntrB-T97P Δ*ntrC* miniTn7[*rhaSR*-PrhaBAD-stRBS-ntrC] (rha-*ntrC*) and Δ*fleQ*Δ*ntrC PFLU1131*-del15 ΔPFLU1132 miniTn7[*rhaSR*-PrhaBAD-stRBS -PFLU1132] (rha-1132). **B)** Motility speed (mm relative to Δ*fleQ ntrB*-T97P) of common pathway ntrB-T97P first step mutant, *PFLU1131*-del15 first-step and PFLU1131/2 promoter second-step mutants. Whiskers represent standard deviation above and below the mean value.

This allows us to conclude that increasing expression of *ntrC* and *PFLU1132* in the presence of their first-step kinase mutations granted stronger motility phenotypes, likely through stronger promiscuous action. The expression level of a transcription factor can therefore significantly impact its propensity for promiscuous regulatory activity, and more highly expressed transcription factor may be expected to engage in non-canonical regulatory interactions more readily.

## Discussion

Essential to understanding why some organisms adapt faster than others in the face of environmental challenges is knowledge of how GRNs and regulatory systems evolve. Evolutionary innovation of transcription factors within these networks depends on gain of functional promiscuity as a first step, however the rules of when and where such properties develop within the context of the GRN is currently unknown. Using our microbial model system, we have identified two available pathways for rescuing flagellar motility in a maladapted Δ*fleQ* network; through mutations to the NtrBC and PFLU1131/2 two component systems. However, one pathway – the NtrBC pathway – is always used to the exclusion of the other when both are present. By comparing these available evolutionary pathways, we can resolve why NtrC is more accessible for gaining promiscuity and rewiring due to its position within its GRN. In addition, we can identify three key properties that facilitate transcription factor promiscuity and opportunities for evolutionary innovation.

**Firstly, our results confirm the importance of shared structural homology between the absent and recruited transcription factor.** In our system, the shared structural homology of NtrC and PFLU1132 with FleQ grants a basal affinity for regulating flagellar genes that evolution can act upon. This likely explains why structurally unrelated transcription factors do not rewire in our model, and indicates that similarity to FleQ facilitates the gain of promiscuity observed in NtrC and PFLU1132. However it is not the case that the regulators that are most structurally homologous to FleQ are the first to be repurposed^32^, so homology is not the deciding factor in which RpoN-EBPs become promiscuous to rescue flagellar motility. It is possible that the RpoN-EBPs represent a hyper-evolvable family of transcription factors through the nature of their connectivity and mechanism of transcriptional regulation via RpoN. To investigate this evolutionary rescue of a non-RpoN dependent trait would need to be tested using a different maladapted GRN rescue model. **Secondly, hyperactivation of the co-opted transcription factor benefits promiscuous regulatory activity**. In our model system, first step mutations that granted a slow motile phenotype were always achieved through mutation of a kinase – resulting in hyper-activation of their cognate transcription factors. Most transcription factors are subject to some form of post-translational control, most commonly through phosphorylation of a REC receiver domain in bacteria^42^.

Promiscuous binding may occur transiently or only briefly, so ensuring that the transcription factor is constantly activated increases the likelihood of these non-cognate regulatory events leading to altered gene expression. **Finally - and perhaps most importantly - high gene expression of the transcription factor boosts promiscuous activity**. High expression of a transcription factor can lead to saturation of native binding sites. Any unbound, active transcription factor is then free to engage in promiscuous activity. These low affinity interactions will occur more readily if there is a higher concentration of transcription factor present, following laws of mass action kinetics^43^. Typically, transcription factors are expressed at low levels in bacteria^44^, due to only a small protein copy number being needed for a small number of binding sites in most cases^45^. Transcription factor expression level may well be optimised at a low level to prevent mis-regulation. Indeed, highly expressed and connected transcription factors were found to result in greater network perturbations when synthetically rewired^15^, which may reflect a greater ability to act promiscuously. Many molecular biology studies use overexpression/overactivation of proteins of interest as a means to determining their biological functions^46^. Our findings also suggest that this approach should perhaps be used more cautiously, certainly when studying transcription factor function.

Aside from mutations directly targeting activation and expression of *PFLU1131/2*, we observed several distinct strategies that enhance PFLU1132 promiscuity through global regulator changes. These strategies likely impact the wider GRN and transcriptome significantly, with mutations to RpoC, RpsK and Rho being particularly striking. RpoC and RpsK mutations may act to increase expression of most genes including *PFLU1131/2* and the flagellar genes. Mutations to the *PFLU1583/4* anti-sigma factor system also indicated a strategy to boost promiscuity that would have a significant impact of the wider GRN. Global regulators are often sites of mutation and innovation^47, 48^, with mutations affecting sigma factor function in particular having large impacts on global gene expression which can drive phenotypic innovation^49^.

Changes to sigma factors have been shown to facilitate GRN remodelling, as was the case for a study of *P. aeruginosa* during adaptation to the CF lung^13^. Our study highlights for the first time that such mutational strategies can also achieve increases in promiscuity for specific transcription factors to facilitate evolutionary innovation.

Each of these mutational strategies that enhanced functional promiscuity in PFLU1132 acted to alter the pre-existing GRN architecture around the transcription factor, whether at a local level by altering the *PFLU1131/2* promoter or at a global regulatory level.

Our findings suggest that pre-existing GRN architecture can be a key constraining factor in gaining functional promiscuity and evolutionary innovation by a transcription factor. For example, the ease by which a transcription factor can gain high expression depends on the GRN structure it sits within. Many transcription factors promote their own expression^50^ (positive autoregulation) and can generate high self-expression through runaway positive feedback. Such positive feedback loops are typically contained by a negative repressor^51–53^, including in the case of NtrBC (Fig.1B). Mutational loss of repression can therefore be an evolutionarily accessible mechanism to achieve promiscuity-enhancing high expression levels of a transcription factor. In our model system, GlnK prevents runaway *ntrBC* autoregulation by inhibiting NtrB^54^ (Fig.1B). All previously observed NtrBC pathway mutations were single *de-novo* mutations that act to remove repression through GlnK, both directly or indirectly^31, 55, 56^. These have a dual effect, both hyperactivating the NtrBC system, and strongly upregulating *ntrBC* expression through their positive autoregulatory loop. In contrast, we have identified in this study that to achieve the same effect of high activity and high expression in PFLU1131/2, multiple mutations were required (Fig.1C). This highlights the clear evolutionary advantage NtrBC has in becoming promiscuous compared to PFLU1131/2.

From the evidence outlined, pre-existing GRN motifs can be predicted to significantly bias the evolvability of the transcription factors under their control. This is strongly supported by the literature: mutational loss of a negative repressor leading to overactivity and overexpression of a positively autoregulated transcription factor has been observed to drive adaptation in several experimental and clinical settings^57–59^. Considering the alternative scenario, a transcription factor that represses its own expression (negative autoregulation) will hinder the gain of a high expression level, due to the nature of self-repression. Interestingly, the hierarchy favouring NtrBC over PFLU1132 matches previous studies of evolutionary pathway heirarchies^33^, where mutational target size predicts that loss of repressors will be favoured over other precise mutations. Our study highlights that in the context of GRN structures, connectivity to other network components can constrain innovation through gain of promiscuity, alongside other effects including mutational target size.

Our study provides empirical evidence for mechanisms and evolutionary drivers of regulatory evolution. GRNs are key to the adaptability of organisms, and our results reveal the genetic regulatory factors that generate evolvable control systems. We find that feedback loops that can generate high expression of regulators after a small number of mutations can facilitate rapid innovation and adaptation. Previously, we have hypothesised that some transcription factors may be ‘primed’ for evolutionary innovation by virtue of the GRN architecture they exist within^26^. Our data support this by suggesting that promiscuity can be more easily evolved in transcription factors with positive feedback structures. These findings can also be utilised in the design of genetic regulatory systems to have reduced evolvability, a key future goal for synthetic biology projects where the evolutionary change of a constructed system is detrimental^60, 61^. Designing systems that avoid easily accessible routes to hyperactivation and hyperexpression of their components could help prevent rewiring of engineered GRNs. Pre-existing GRN structure can be expected to significantly bias patterns of diversification in regulatory systems, and subsequently shape genomic evolution by influencing the speed in which genetic circuitry can adapt to changing conditions. This is highly important for populations adapting during sudden changes such as niche transitions including the emergence of new pathogens. Our data moves us closer to the eventual goal of empirically building a set of principles by which GRNs adapt and offers unique insights into how these systems can function and influence the evolution of their components.

## Materials and Methods

### Strains and culture conditions

All ancestral lines in this study are derived from *Pseudomonas fluorescens* SBW25Δ*fleQ* IS-ΩKm-hah: PFLU2552 (referred to throughout as *ΔfleQ*). Removal of the flagellum master regulator FleQ and transposon-insertional disruption of the gene *viscB* (PFLU2552), rendering this strain immotile as detailed previously^31, 62^. All strains were cultured on lysogeny broth (LB; Miller) media at 27°C. *Escherichia coli* strains for cloning were cultured on LB media at 37°C. Strains, plasmids, media supplements and primers are detailed in Supplementary Excel file E4.

### Knockout mutant construction by two-step allelic exchange

Knockouts of ntrC and PFLU1132 were achieved using two-step allelic exchange following the protocol of Hmelo et al., 2015 with some alterations. In brief, 400bp flanking regions up- and down-stream of the target gene were amplified and joined by strand overlap extension (SOE) PCR to create a knockout allele. This was inserted into the allelic-exchange suicide vector pTS1 (Containing *sacB* and *tetR*) by SOE-cloning^64^ to create the knockout plasmid, and transformed into the conjugal *E. coli* strain ST18 by chemical-competence heat-shock. Knockout plasmids were transferred from *E. coli* ST18 to *ΔfleQ* by two-parent puddle-mating and merodiploids selected for on LB supplemented with kanamycin sulphate and tetracycline hydrochloride but lacking 5-ALA supplement required for growth of the *E. coli* ST18 auxotrophic Δ*hemA* mutant. Merodiploids were cultured overnight in LB broth lacking tetracycline selection and diluted before spread plating onto NSLB media supplemented with 15% w/v sucrose for *sacB* levansucrase-mediated counterselection of the pTS1 plasmid backbone. Sucrose-resistant colonies were isolated and screened for tetracycline sensitivity. Chromosomal presence of the knockout allele and absence of target gene coding sequences was confirmed by colony PCR. Knockout of ntrC left an ORF encoding an 11 amino acid truncated protein, and knockout of PFLU1132 left an ORF encoding an 8 amino acid truncated protein. All other genomic features including operon structure and terminator regions were left intact.

### Motility evolution experiments

*ΔfleQ*Δ*ntrC* and *ΔfleQ*Δ*ntrC*Δ*PFLU1132* were challenged to rescue motility in the absence of the FleQ master flagellar regulator on soft agar, as described previously^31, 56^. Pure colonies were picked and inoculated into 0.25% agar LB plates made as described in Alsohim et al., 2014, and incubated at 27°C. Plates were checked a minimum of twice daily for motility, recording time to emergence. Motile zones were sampled immediately and always from the leading edge. Motile isolates were streaked on LB agar, and a pure colony picked and stored at -80°C as glycerol stocks of LB overnight cultures. All subsequent analysis was conducted on these pure motile isolates. Experiment was run for six weeks and any replicates without motility after this cut-off recorded as having not evolved.

### Bacterial growth and motility fitness assays

The fitness of the motility phenotype of evolved *ΔfleQ*Δ*ntrC* strains was tested by measuring distance moved over 24 hrs of incubation in 0.25% agar LB plates. Six biological replicates of each strain were grown as separate overnight cultures.

Cultures were adjusted to an OD595 = 1 and resuspended in PBS. Soft agar plates were inoculated with 1 μL of these suspensions, by piercing the surface of the plate with the pipette tip, and then effusing the sample into the gap left by the tip. Plates were incubated for 24 h at 27°C, and photographs taken of motile zones. Surface area moved was then calculated from the radius of the concentric motile zone measured from these images (A = π r²). Values were square root transformed before plotting.

Growth in shaking LB broth was measured by inoculating 99 μL of sterile LB broth with 1 μL of the OD595=1 PBS cell suspensions for each replicate in a 96-well plate. Plates were incubated at 27°C with 180 rpm shaking in a plate reader, recording OD595 every hour for 24 hours. Area under the bacterial growth curve was calculated using the growthcurver package in R and plotted^65^. Area under the bacterial growth curve provides a metric for fitness, as it accounts for the characteristics of lag- and log-phase growth, as well as the final carrying capacity of the population. All growth curves were plotted and inspected prior to calculation of AUGC to ensure similar AUGC values corresponded to similar curve shapes (Supplementary Fig. S3)

### Mutation identification by whole genome resequencing and PCR amplicon sanger sequencing

To identify motility rescuing mutations, genomic DNA was extracted from motile strains and their ancestral strain using the Thermo Scientific GeneJET Genomic DNA Purification Kit. Genomic DNA was quality checked using BR dsDNA Qubit spectrophotometry to determine concentration and nanodrop spectrophotometry to determine purity. Illumina HiSeq sequencing was provided by MicrobesNG (Birmingham, UK), with a minimum 30x coverage and a sample average of 114x coverage. Returned paired-end reads were aligned to the *P. fluorescens* SBW25 reference genome^37^ using the Galaxy platform^66^. INDELs were identified using the integrated genomics viewer^67^. For SNP identification, the variant calling software SNIPPY was used with default parameters (Seemann, 2015). Protein domains affected by mutations detailed in this article were predicted from amino acid sequences using pfam, SMART and BLASTp. For a subset of additional motile mutants, PCR amplification and subsequent sanger sequencing of *PFLU1131* was performed as this gene was mutated in all previously sequenced strains. This was done using the service provided by Eurofins Genomics. PCR amplicons were purified using the Monarch® PCR & DNA Cleanup Kit (New England Biolabs).

Mutations were identified by alignment of the returned PFLU1131 sequence against the *P. fluorescens* SBW25 reference genome using NCBI BLAST.

Three second-step isolates derived from line 24 were sampled due to observed colony morphology variation. After whole genome re-sequencing it became apparent that this variation was due to the presence of a *mutS* mutation and not due to diverse motility strategies. These isolates all have the mutation *PFLU1132*-D111G (Supplementary excel file E1, isolate ID’s 24-S1, 24-LCV, and 24-SCV), however this is likely not due to parallel evolution, but instead a single instance of a *PFLU1132*- D111G mutant that then diversified due to *mutS* mutation. For all subsequent analysis these three isolates were treated as a single instance of *PFLU1132*-D111G.

### RNA-sequencing

Whole-cell RNA was extracted from 20 OD units of *P. fluorescens* cultures in mid-log phase growth (OD595 ∼1.5). Cultures were incubated in LB broth at 27°C and 180 rpm shaking. Extractions were performed for biological triplicates of each strain of interest. Upon reaching the desired OD, growth and RNA expression was halted by addition of a ½ culture volume of ice-cold killing buffer (20 mM NaN3, 20 mM Tris-HCl, 5 mM MgCl2). Cells were pelleted, and the killing buffer removed. A lysis buffer of β-mercaptoethanol in buffer RLT from the Qiagen RNeasy extraction kit was used to resuspend pellets, which were then lysed by bead-milling at 4500 rpm for 45 s with lysing matrix B. Lysates were spun through columns from the Qiagen RNeasy extraction kit, and the extraction completed following the RNeasy kit protocol. A DNase I treatment step was included between washes with buffer RW1, by adding DNase I directly to the column from the RNase-free DNase kit (Qiagen) following kit protocol. Samples were eluted in nuclease-free water, and subsequently treated with TURBO DNase from the Turbo DNA-free kit (Invitrogen) following kit protocols.

Purified RNA concentration was measured by Qubit RNA BR assay (Thermo- scientific), RNA quality by nanodrop spectrophotometry, and RNA integrity by agarose gel electrophoresis. RNA sequencing was provided by Oxford Genomics (Oxford UK). Samples were ribodepleted, and the mRNA fraction converted to cDNA with dUTP incorporated during second strand synthesis. The cDNA was end-repaired, A-tailed and adapter-ligated. Prior to amplification, samples underwent uridine digestion. The prepared libraries were size selected, multiplexed and QC’ed before paired end sequencing over one unit of a flow cell in a NovaSeq 6000.

Returned data was quality checked before it was returned in fastq format. Reference-based transcript assembly was performed using the Rockhopper RNA sequencing analysis software^68^, to generate transcript read counts. Differential gene expression analysis was performed using the DESeq2 package of the R statistical coding language to create lists of differentially expressed genes and corresponding Log2Fold changes in gene expression.

### Rhamnose-inducible expression system constructs

Complementation of *ntrC* and *PFLU1132* knockouts, as well as inducible expression experiments were performed by single-copy chromosomal re-introduction of the RpoN-EBP genes under control of a *rhaSR-PrhaBAD* L-rhamnose inducible expression system^69^. The RpoN-EBP ORF was amplified and a strong ribosome binding site (stRBS) introduced upstream after a 7bp short spacer between the site and the start codon. This construct was placed downstream of the *PrhaBAD* promoter on the miniTn7 suicide-vector pJM220 (obtained from the Addgene plasmid repository, plasmid #110559) by restriction-ligation and transferred to *E. coli* DH5α by chemical-competence heat-shock. The miniTn7 transposon containing the *rhaSR* genes and the PrhaBAD-stRBS-RpoN-EBP construct were transferred to the *P. fluorescens* chromosome by transposonal insertion downstream of the *glmS* gene via four-parent puddle-mating conjugation^70^. The relevant *E. coli* DH5α pJM220- derived plasmid donor was combined with recipient *P. fluorescens ΔfleQ* strains, transposition helper *E. coli* SM10 λpir pTNS2 and conjugation helper *E. coli* SP50 pRK2073, and Gentamicin resistant *Pseudomonas* selected for on LB supplemented with Gentamicin sulphate and Kanamycin sulphate. Chromosomal insertion of the correct miniTn7 transposon was confirmed by colony PCR.

### β-galactosidase activity assay

To validate the expression dynamics of the rhaSR-PrhaBAD system in the genetic backgrounds used in this work, the same system described above was introduced into *P. fluorescens ΔfleQ* and *ΔfleQ*Δ*ntrC* with *lacZ* under the control of PrhaBAD. This was obtained from the pJM230 plasmid^69^ (obtained from the Addgene plasmid repository, plasmid #110560). Overnight cultures of each strain were set up in biological triplicate, and used to inoculate 9 mL cultures of LB broth supplemented with a range of L-rhamnose concentrations at a bacterial OD600 = 0.05. Cultures were incubated shaking 180 r.p.m. at 27°C until they reached and OD600 = 0.5, 2 mL was spun down and frozen at -20°C until needed. Cell pellets were resuspended in Z-buffer (60mM Na2HPO4*7H2O, 40mM NaH2PO4*H2O, 10mM KCl, 1mMMgSO4*7H2O), OD600 recorded, and 400μL cells added to 600μL fresh Z-buffer. Cell suspension was lysed with addition of 40 μL Chloroform, 20 μL 0.1% SDS and vortexing followed by 10 minutes incubation at 30°C. β-galactosidase activity assay was begun with addition of 200μL o-nitrophenyl-β-D-galactopyraniside (ONPG) and incubated at 30°C for 20 minutes. If reactions become visibly yellow they were halted immediately by addition of 500 μL Na2CO3 and the time recorded. All other reactions were halted in this manner after 20 minutes. Reactions were centrifuged to remove debris and 900 μL added to a cuvette and A420 measured. LacZ reporter activity (MU) was calculated by: MU = (A420 * 1000) / (Time(mins) * 0.4 * OD600 of initial Z-buffer cell suspension).

### Statistical analyses and data handling

All statistical analysis and data handling was performed using R core statistical packages and the Dunn.test package. Shapiro-Wilks normality tests were performed to confirm non-normality of datasets. To compare group medians for more than two groups, a Kruskal-Wallis test with post-hoc Dunn test and Benjamini-Hochberg correction was performed, with a P ≤ 0.025 taken to indicate significance. For comparisons of two groups only, a two-sample Wilcox test was used. Spearman’s correlation was used to test significance of correlations between motility rate and rhamnose concentration, using the R core statistical function cor.test with the spearman method specified.

For analysis of transcriptomic data, RNA sequencing output was run through Rockhopper for reference-based transcript assembly. Transcript count data were then analysed using DESeq2 package in R, for differential gene expression analysis, as well as generation of PCA and Volcano plots. A gene was taken as being differentially expressed if it differed by a Log2Fold change of ≥1 or ≤ -1 between two samples (representing a doubling or halving of expression level respectively) with an adjusted P-value of ≤0.001.

To identify whether Rho dependent terminators were present downstream of genes of interest, the RhoTermPredict algorithm^71^ was run on the *P. fluorescens* SBW25 reference genome in Python.

## Supporting information

Supplementary excel sheet E1

Supplementary excel sheet E2

Supplementary excel sheet E3

Supplementary excel sheet E4

Supplementary excel sheet E5

---

**Supplementary Fig S1:**
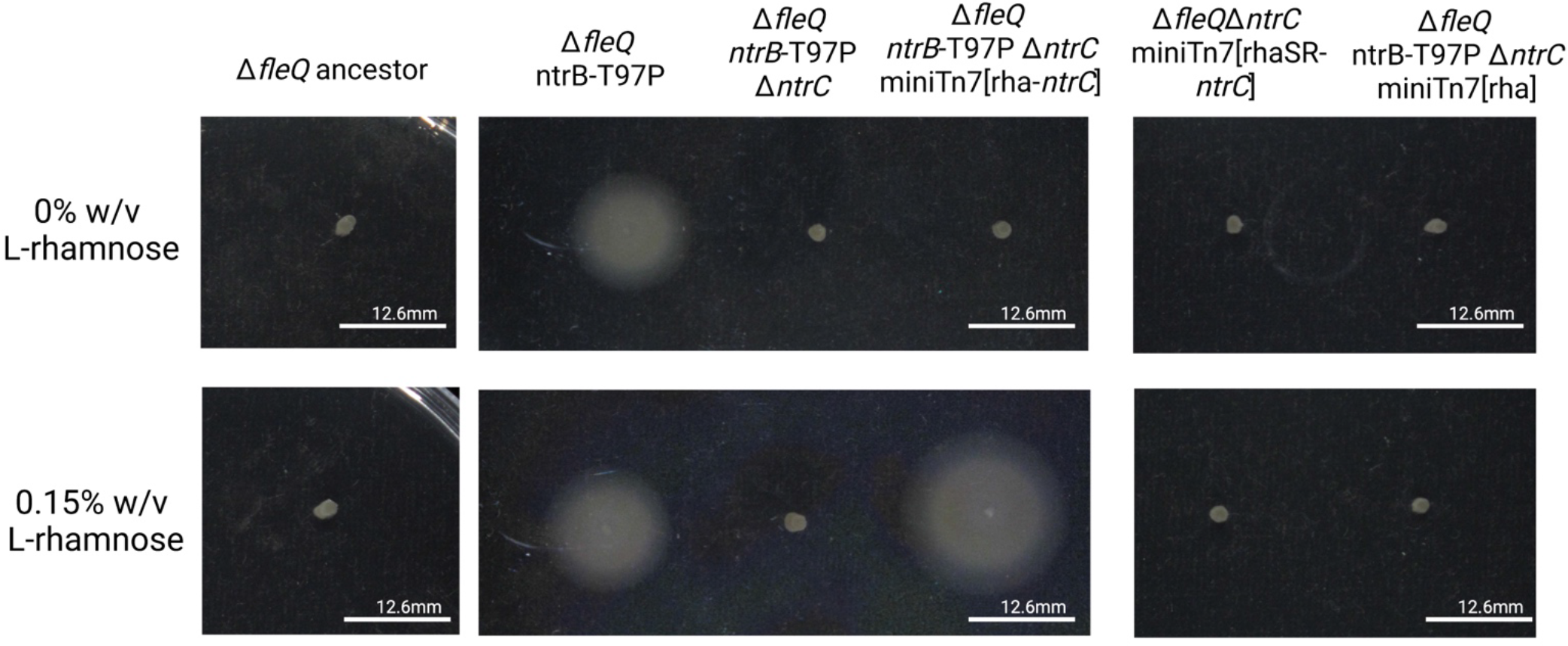
FleQ-homolog encoding gene *ntrC* is essential for rescued flagellar motility in *ntrB* mutant strain and depends on the presence of a kinase mutation. Transcription factor gene NtrC was deleted, and then reintroduced as a single-copy chromosomal insertion expressed from an L-rhamnose inducible promoter system (*rhaSR-PrhaBAD*). The same complementation lacking the ntrB- T97P mutation, as well as an empty expression system transposon were included as further controls. Photographs of motility after 1 day incubation in 0.25% agar LB plates supplemented with or without 0.15% L-rhamnose for induction of transcription factor expression.

**Supplementary Figure S2.**
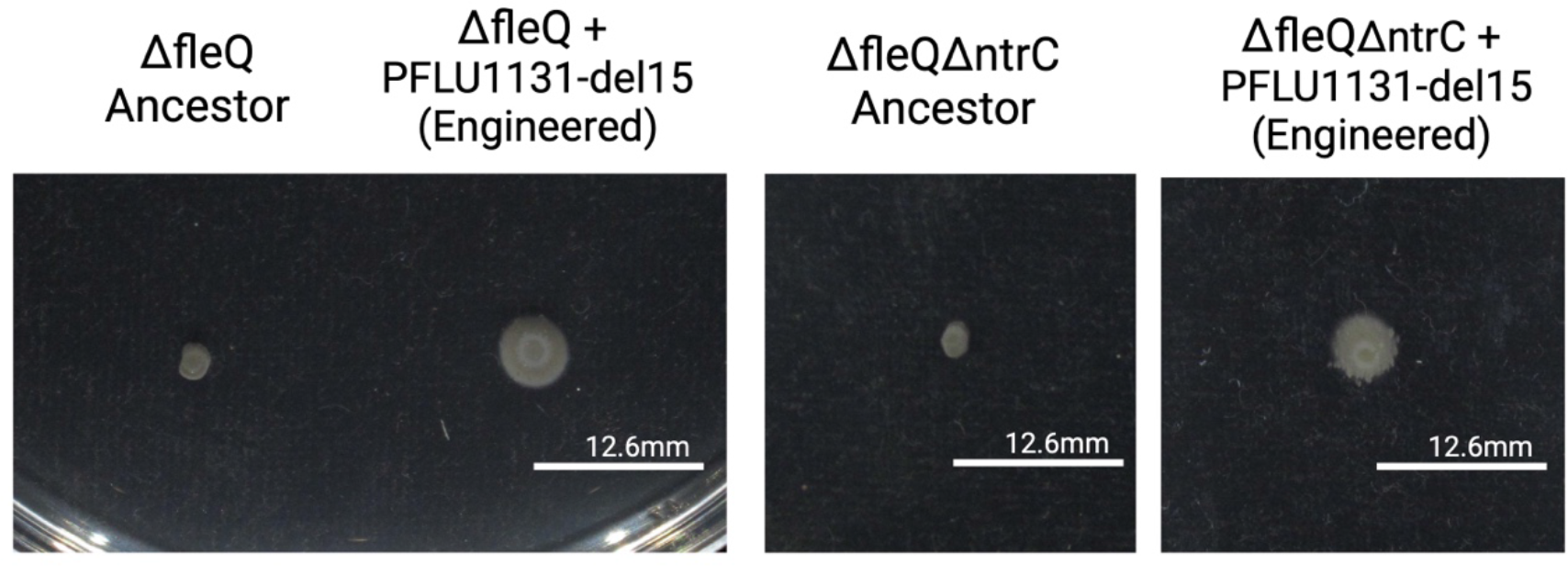
: PFLU1131-del15 mutation engineered into the Δ*fleQ* and Δ*fleQ*Δ*ntrC* ancestral lines provides motility in both conditions. Indicates that rescue of motility by this mutation does not depend on Δ*ntrC* and is the sole mutation required to do so.

**Supplementary Figure S3:**
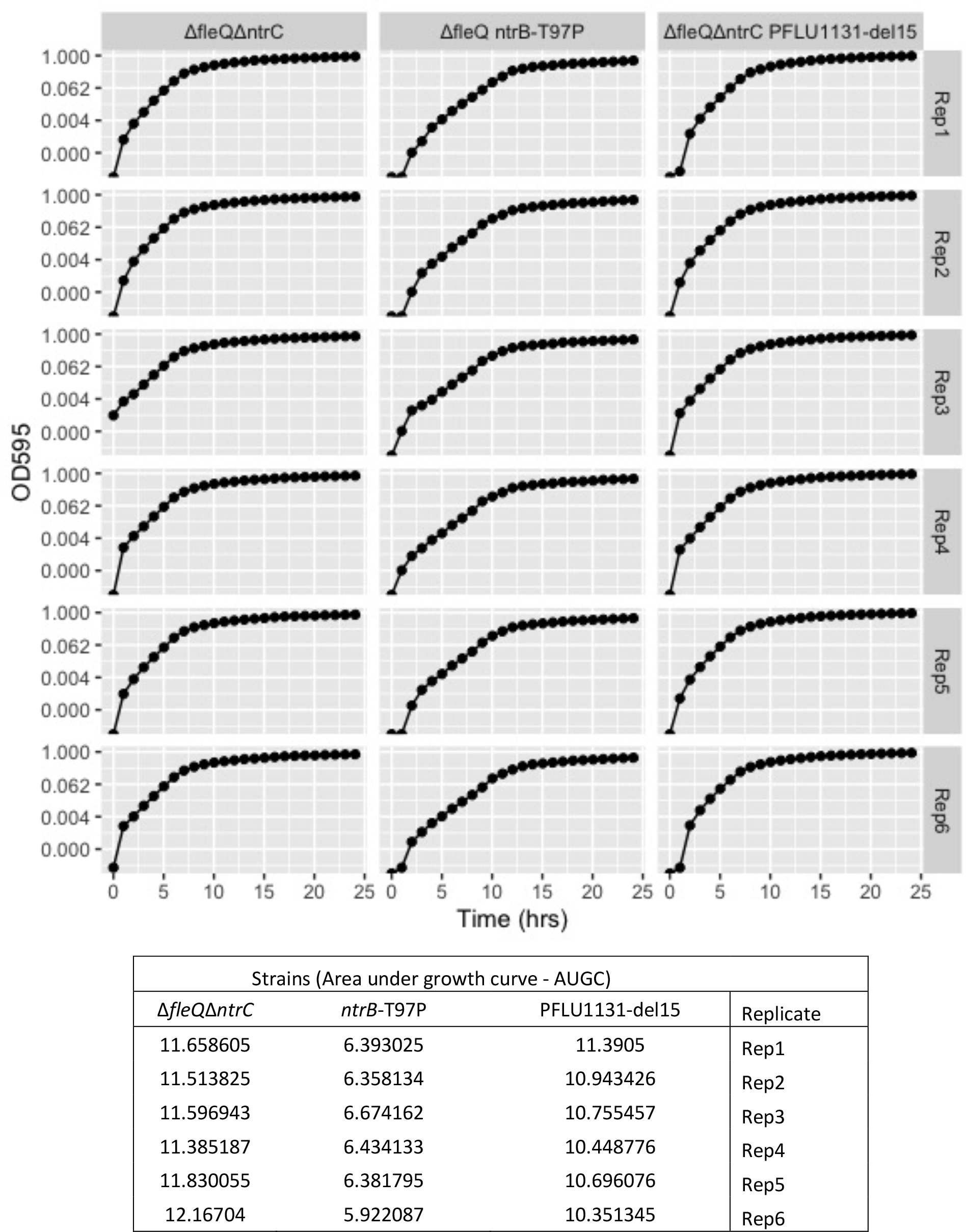
Individual microbial growth curves that provide area under the growth curve (AUGC) values that are plotted in Fig.4C. Six biological replicates for each of the three strains were assayed for change in OD595 over 24 hours of incubation in shaking LB broth. AUGC values corresponding to each curve are displayed in the table below the graphs.

**Supplementary Figure S4.**
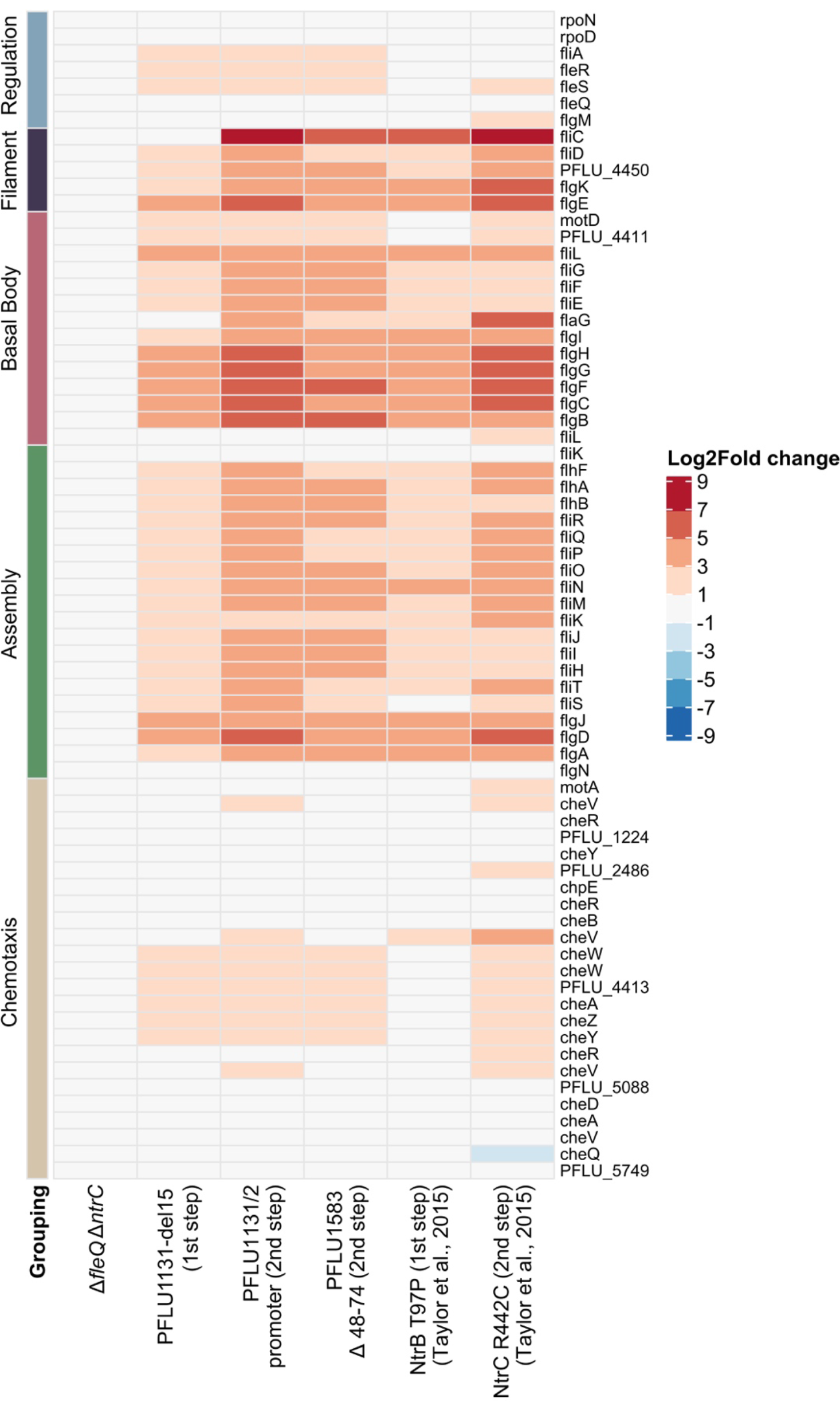
: Impact of motility rescue mutations of flagellar and chemotaxis gene expression. Log2Fold changes in gene expression relative to the *ΔfleQ* ancestor are shown for all genes associated with flagellar motility in *P. fluorescens* SBW25. Functional groups of genes are indicated by the coloured bars and labels to the left of the plot.

**Supplementary Figure S5.**
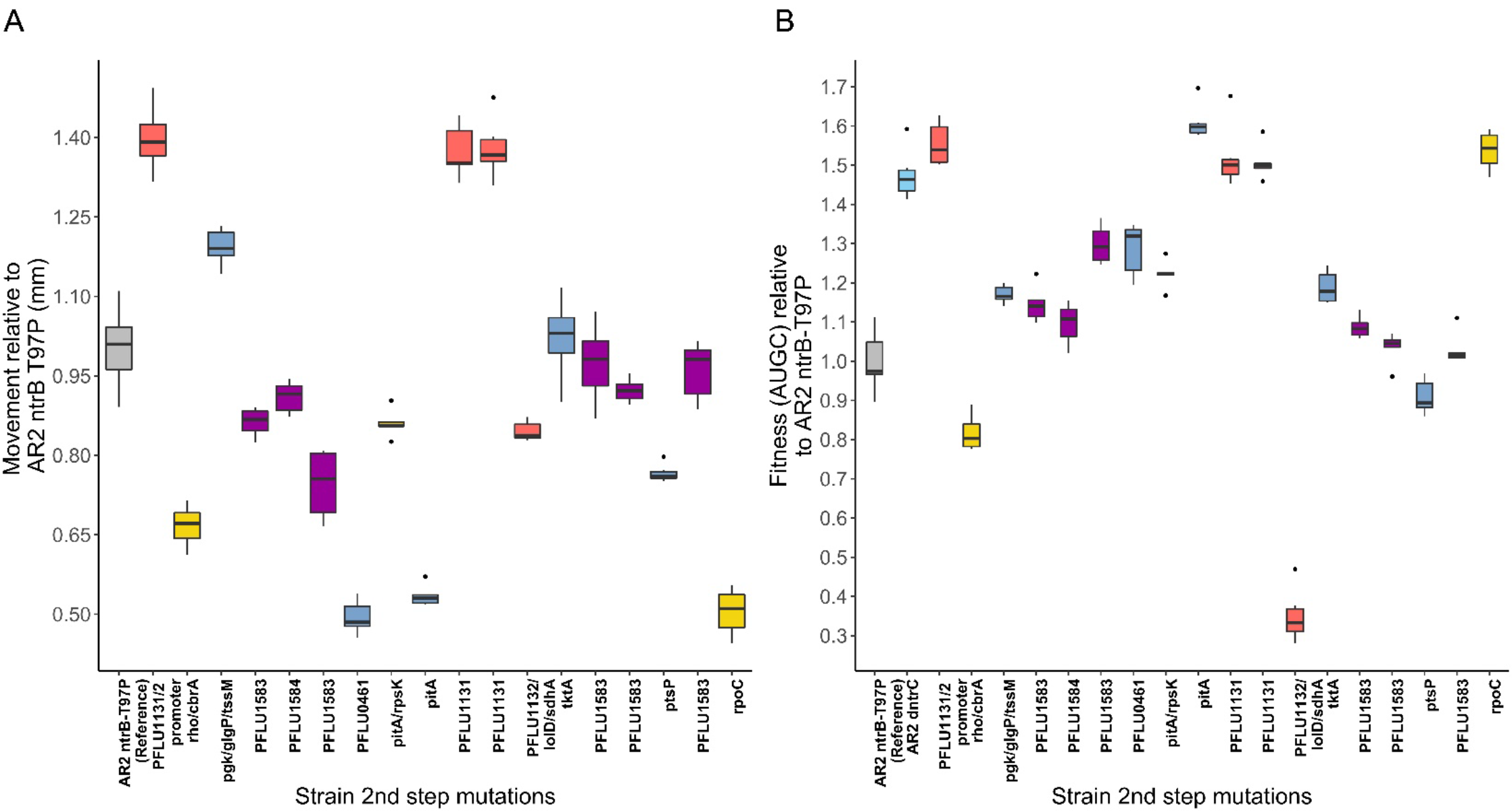
: Phenotypes of motility rescue second-step mutants. **A)** Race assay as measure of motility fitness. Distance moved over 24hrs in 0.25% Agar LB plates measured relative to the *ΔfleQ* ancestor *ntrB*-T97P mutant. **B)** Fitness in LB broth measured as area under the 24hr growth curve (AUGC) relative to *ΔfleQ* ancestor *ntrB*-T97P. In both plots, mutations are coloured by functional category: Red – PFLU1131/2, Purple – PFLU1583/4, Blue – Metabolic, Yellow – Global regulatory, Grey – other. For all boxplots – box represents first to third quartile range, middle line represents median value, whiskers range from quartiles to maxima and minima.

**Supplementary Figure S6.**
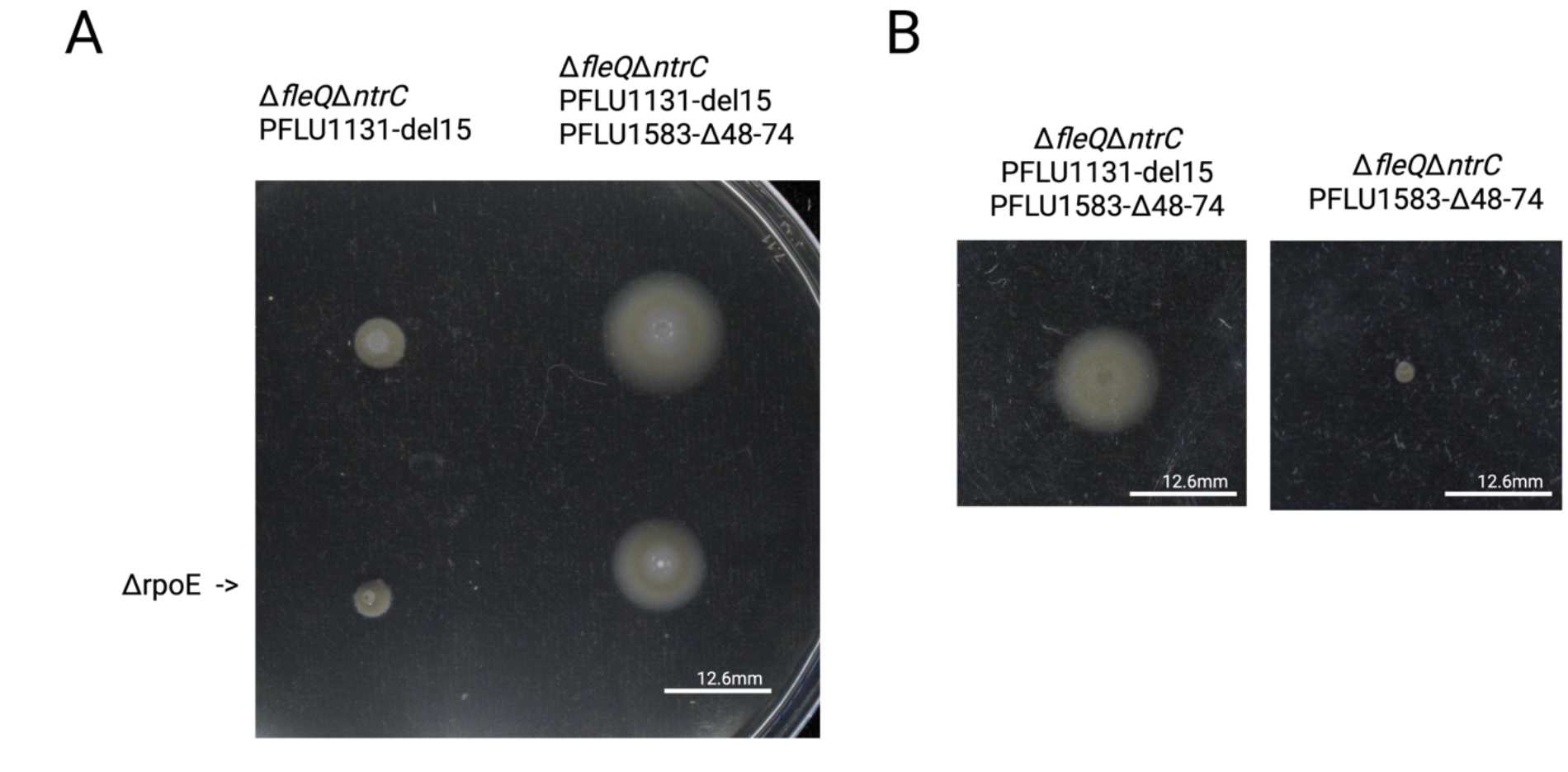
: PFLU1583 Δ48-74 mutation depends to presence of PFLU1131 mutation, and does not act through sigma factor RpoE. **A)** Motility of PFLU PFLU1583 Δ48-74 with and without accompanying *PFLU1131*-del15 mutation. **B)** *rpoE* knockout does not revert PFLU1583 mutant to first-step motility phenotype.

**Supplementary Figure S7.**
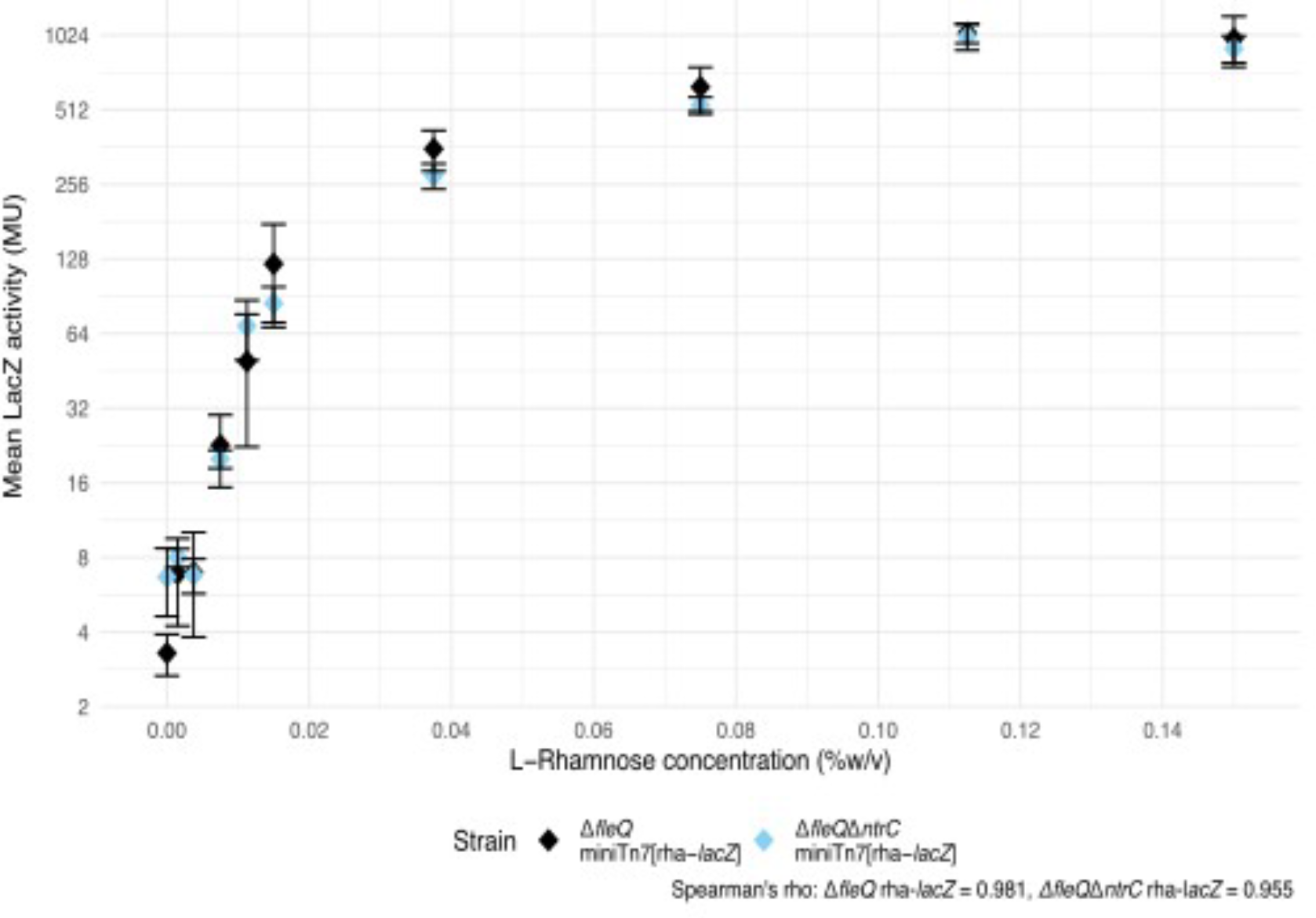
: Activity of LacZ expression reporter (Miller units – MU) with increasing L-rhamnose concentration (%w/v) in both the *ΔfleQ* and *ΔfleQΔntrC* ancestral genetic backgrounds with miniTn7[rhaSR-PrhaBAD-stRBS-LacZ]. Whiskers represent standard deviation above and below the mean value.

## Notes

https://osf.io/pcdhx/

### Competing Interest Statement

The authors have declared no competing interest.

### Summary of Updates

Figures and main text significantly revised to improve clarity and presentation of results.

